# Microbial mat compositions and localization patterns explain the virulence of black band disease in corals

**DOI:** 10.1101/2022.10.04.510905

**Authors:** Naohisa Wada, Akira Iguchi, Yuta Urabe, Yuki Yoshioka, Natsumi Abe, Kazuki Takase, Shuji Hayashi, Sakiko Kawanabe, Yui Sato, Sen-Lin Tang, Nobuhiro Mano

**Affiliations:** Biodiversity Research Center, Academia Sinica, No.128, Sec 2, Academia Rd, Nangang, Taipei 11529 Taiwan; Department of Marine Science and Resources, College of Bioresource Science, Nihon University, Fujisawa, Kanagawa, Japan; Geological Survey of Japan, National Institute of Advanced Industrial Science and Technology (AIST), 1-1-1 Higashi, Tsukuba, Ibaraki 305-8567, Japan; Research Laboratory on Environmentally-conscious Developments and Technologies [E-code], National Institute of Advanced Industrial Science and Technology (AIST), Tsukuba 305-8567, Japan; Department of Bioresources Engineering, National Institute of Technology, Okinawa College, 905 Henoko, Nago-City, Okinawa 905-2192, Japan; College of Science and Engineering, James Cook University, Townsville, Queensland, Australia

**Keywords:** Coral disease, Black Band Disease, Migration, Progression, *Arcobacteraceae*

## Abstract

Black band disease (BBD) in corals is characterized by a distinctive, band-like microbial mat, which spreads across the tissues and often kills infected colonies. The microbial mat is dominated by cyanobacteria but also commonly contains sulfide-oxidizing bacteria (SOB), sulfate-reducing bacteria (SRB), and other microbes. The migration rate in BBD varies across different environmental conditions including temperature, light, and pH. However, whether variations in the migration rates reflect differences in the microbial consortium within the BBD mat remains unknown. Here, we show that the micro-scale surface structure, bacterial composition, and spatial distribution differed across BBD lesions with different migration rates. The migration rate was positively correlated with the relative abundance of potential SOBs belonging to *Arcobacteraceae* localized in the middle layer within the mat and negatively correlated with the relative abundance of other potential SOBs belonging to *Rhodobacteraceae*. Our study highlights the microbial composition in BBD as an important determinant of virulence.

## Introduction

Cyanobacteria are often key organisms that form microbial mats in natural environments. A striking feature of the cyanobacterial mat is its stratified structure and specific layers in which different trophic microorganisms are distributed^1^. The uppermost layers are generally dominated by aerobic cyanobacteria, diatoms, and other oxygenic phototrophs, while the lowest layers are dominated by various anaerobic bacteria. The cyanobacterial mats occur in terrestrial and aquatic environments such as tidal sand flats, hypersaline ponds, hot springs, and intertidal zones^1,2^. The vertical distributions of bacteria can fluctuate daily^3^, and the mats can expand horizontally in radial directions^4^.

Coral black band disease (BBD), characterized by a dark, cyanobacterial-dominated microbial mat, exhibits a unique band shape that linearly migrates through living coral tissues (**Fig. 1**). The characteristic black band migrates over living coral tissues, resulting in lysis and necrosis of the underlying tissue, and leaving behind a bare coral skeleton^5^. The widths of the black band between apparently normal coral tissue and freshly exposed skeleton can range from a few millimeters to seven centimeters^6,7^. The migration rate of BBD, recorded up to 2 cm/day^8^, vary with temperature^9,10^, light^9,10^, pH^11^, and different geographical conditions^12^. As many as five-fold differences in the migration rate concurrently occurred within a single reef^13^.

**Fig. 1.**
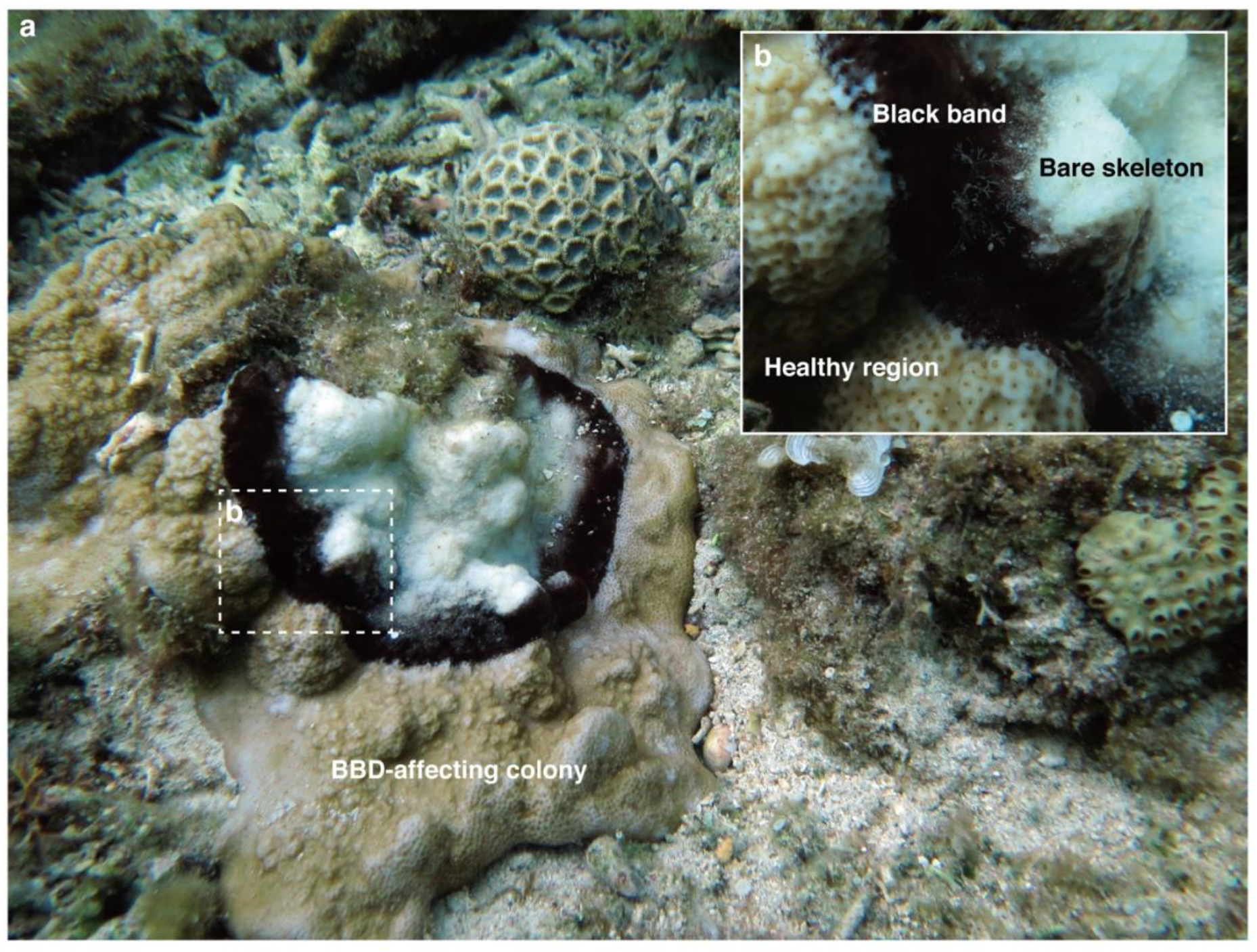
A coral colony *Montipora* sp. being infected by black band disease (BBD) in Aka Island, Okinawa, Japan. **(a)**. Close-up of the interface showing black band, which was composed of cyanobacterial mat between live coral tissue (healthy region) and exposed coral white skeleton (bare skeleton) (**b**).

The biomass of the polymicrobial mats is dominated by the filamentous cyanobacterial genera *Oscillatoria, Roseofilum*, and *Pseudoscillatoria* that had been identified from the Indo-Pacific, Caribbean, and Red Sea, respectively^14^. Those cyanobacteria belong to a monophyletic lineage and are consistently found in BBD lesions, indicating their pivotal role in the BBD etiology^14^. Other common microbial constituents of the BBD lesion are SOB (e.g., *Beggiatoa* spp.^15^ and *Rhodobacterales*^16^), SRB (e.g., *Desulfovibrio*^17^ and *Desulfobacteraceae*^18^), some diverse heterotrophic bacteria^14^, and archaea^19^. Cyanobacteria are located in the upper part of the microbial mat; SOB, SRB and heterotrophic bacteria are found in the lower part during the daytime; and the aerobic bacteria, including SOB, move over the cyanobacterial layer at night^14^. A mechanism for this special distribution has been proposed. The cyanobacteria’s activity during the daytime results in a dynamic vertical micro-gradient of oxygen^7,20^. During the night, an oxygen-minimal environment is formed within the mat, and then the SRB proliferate and activate in the anoxic conditions underneath the mat^7^. The oxygen deprivation and high concentrations of sulfide from SRB are fatal to coral tissue, and thus are considered the most important factors in the etiology of BBD^14,21^. Toxins of cyanobacteria have also been implicated in coral tissue necrosis at the lesion-front in some studies^22^. Meanwhile, SOB members in BBD are inexplicable so far. Common SOB members (such as *Beggiatoa* spp. and/or *Rhodobacterales*) represent a very minor component of the BBD microbial consortia that were obtained from widely different geographic locations^16,18,19,23–27^. This suggests that the SOB activity itself is not the primary driver of BBD pathogenesis, but that its scarcity may aid in the accumulation of sulfide within the BBD lesions^14^. Nonetheless, other research from the Red Sea reported that an *Arcobacter* sp. (i.e., the other viable SOB members) was detected highly abundant in BBD lesions^27^. Considering the complexity of the BBD microbial consortium, the etiology and underlying mechanisms of pathogenesis and virulence of BBD still remain unresolved^14^, especially when it comes to their connection to bacterial composition and localization. In addition, fine-scale microbial localizations within BBD lesions have not been entirely explicated, and doing so could elucidate the BBD-polymicrobial dynamics^14^.

To characterize the microbial consortium involved in the migration rate of BBD and pathogenesis, we examined whether the fastest migration rate of BBD reflects an ideal bacterial composition with high virulence. This study specifically examined how the polymicrobial consortia within BBD lesions differ across migration rates by using a scanning electron microscope (SEM), bacterial community sequencing, and a combined method of fluorescence *in situ* hybridization (FISH) and undecalcified coral sectioning. To account for regional variations, BBD samples were collected from two geographic locations in Okinawa, Japan.

## Results

### Surface structures of BBD among various migration rates of BBD

First, we measured linear-migration rates in BBD lesions of the encrusting coral *Montipora* spp. in two locations separated by an oceanic distance of approximately 70 km (Sesoko Island and Aka Island in Okinawa, Japan, **Suppl. fig. S1**). Eighteen BBD lesions showing the various linear-migration rates (ranging from 0.30 to 5.96 mm/day) were collected from these two locations. SEM images of the BBD mat surface adjacent to healthy coral tissue morphologically showed the presence of numerous microbes comprising thinner filamentous cyanobacteria (**Suppl. fig. S2a**), relatively thicker filamentous cyanobacteria (**Suppl. fig. S2b**), filamentous microorganisms (**Suppl. fig. S2c)**, other rod-shaped bacteria (**Suppl. fig. S2d)**, and two types of ciliates (**Suppl. fig. S2e**-**f**). Those microbes were morphologically similar in the samples from the two locations. Regardless of the migration rate of the lesions, structures of extracellular polymeric substances (EPS) were observed across the surface of the cyanobacterial aggregation in the samples from Aka Island, whereas structures of EPS were generally not observed across each of the individual cyanobacterial filaments from Sesoko Island (**Fig. 2**). In the BBD area, within 1 mm of the border of healthy coral tissue, the thinner filamentous cyanobacteria (**Suppl. fig. S2a**) dominated the mat consistently in all the samples, regardless of location and migration rate (**Fig. 2**). The thicker filamentous cyanobacteria (**Suppl. fig. S2b**) were found only in the samples with migration rates of 2.74 mm/day from Aka Island and 4.18 mm/day from Sesoko Island. The filamentous microorganisms (**Suppl. fig. S2c**) appeared in the samples from both locations with migration rates from 1.89 mm/day to 3.99 mm/day. The ciliates displayed two different shapes: an elongated, tube-shaped body (Type A, **Suppl. fig. S2e**) and a pellet-like body (Type B, **Suppl. fig. S2f**). The type A and B ciliates were observed widely on the surface across samples with various migration rates ranging from 0.30 – 5.63 mm/day and 1.53 – 5.96 mm/day, respectively.

**Fig. 2.**
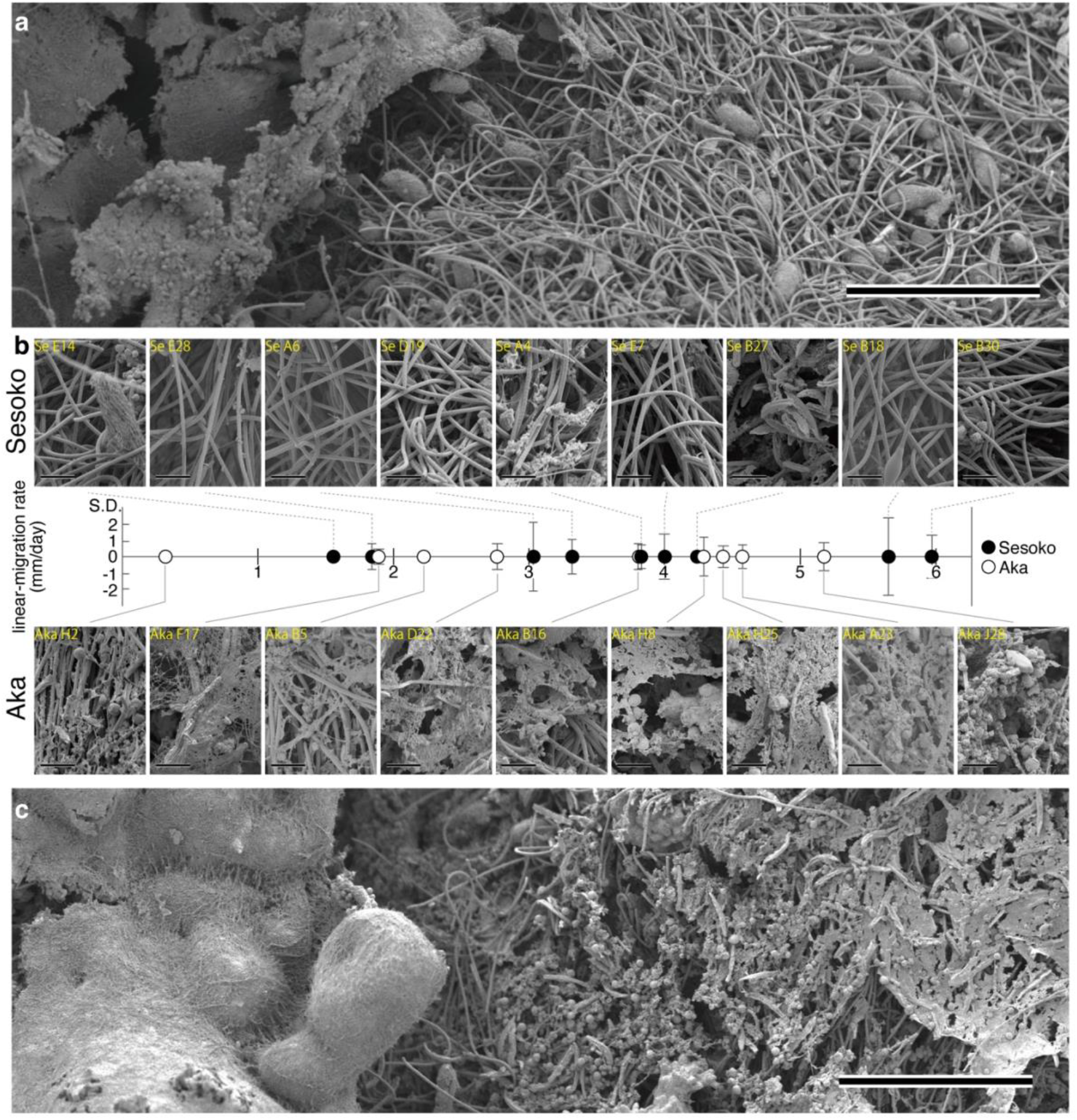
SEM images displaying the surface structures of BBD from two locations with different migration rates. The sample from Sesoko Island (**a**, upper parts in panel **b**) displaying cyanobacterial filaments with non-impurities on the surfaces than Aka Island. In Aka Island, the samples (lower parts in panel **b** and **c**) showed extracellular polymeric substances (EPS) on the surfaces. Comparison of surface appearances between two locations across various migration rates (**b**). Scale bars indicate 200 µm (**a** and **c**) and 20 µm (**b**). The direction of BBD-migration is from right to left (**a** and **c**).

### Bacterial community in the BBD mat

For bacterial community analysis, twelve separate BBD bands, showing different linear-migration rates ranging from 2.13 to 6.36 mm/day (**Fig. 3a**), and a variety of bacterial communities (**Fig. 3b**) were collected from the same two locations mentioned above. The alpha diversity of bacteria (revealed by observed OTUs and Chao1 richness indices) were negatively correlated with migration rates (Spearman’s rank correlation, rho = −0.80, *p* value < 0.01, and rho = −0.74, *p* value < 0.001) (**Fig. 3c**). To avoid biases due to the compositional data in the correlation with migration, we applied a centered log-ratio (clr) transformation on our data for the downstream analyses. Transformation-based principal component analysis (tb-PCA) showed that the locational differences had a negligible effect on the differences in bacterial communities at the family level (**Fig. 3d**). To assess the effect of the migration gradient on the community structure, we tested generalized additive models with an integrated smoothness estimation (R2: 0.135, *p* value: 0.534) and a Mantel test (*R*: 0.06786, *p* value: 0.297). The results were worse at explaining the quantitative relationship between migration rates to the bacterial family structure. However, the pattern of partial correlation between representative bacteria at the family level with migration rates was verified by the clr transformation matrix. Only two potential SOB families, the *Rhodobacteraceae* and *Arcobacteraceae*, were significantly correlated negatively and positively with linear-migration rates (Spearman’s rank correlation, rho = –0.59 *p* < 0.05, rho = 0.85 and *p* < 0.01), respectively (**Fig. 4** and **Suppl. table S3**). 44 OTUs (operational taxonomic unit[s]) in all samples belonged to the family *Rhodobacteraceae* (**Suppl. table S1**), and only two OTUs (OTU 6 and OTU24), ranging from 0.01 to 13.47% and from 0.01 to 6.96%, respectively, had > 1% of total relative abundance (**Suppl. table S2**). OTU 6 was assigned to the genus *Ruegeria*, and exactly matched the sequence of an uncultured alpha proteobacterium 128-64 (AF473938) that was retrieved from BBD in the Caribbean Sea (**Suppl. table S2**). OTU 24 was identified as the genus *Thalassobius* and was 100% similar to both an uncultured bacterium clone BBD-Aug08-3BB-36 (GU472129) and an uncultured bacterium clone Otu0020 (MH341656) in a BBD mat from the Red Sea (**Suppl. table S2**). As the viable sulfide-oxidizers or heterotrophic bacteria (see more detail in discussion), *Arcobacteraceae* were found with 14 OTUs in all BBD samples across the various migration rates, three of which were OTU 8, OTU 9, and OTU11, which ranged between 0.54-10.84% in six samples, 0.03-10.81% in nine samples, and 0.01 - 5.46% in all samples, respectively (**Suppl. table S2**). All three OTUs in the *Arcobacteraceae* were closely related to bacteria (with 98.88 to 100% similarity) that were observed in BBD in the Indo-Pacific Ocean and the Caribbean Sea (**Suppl. table S2**). Six of the 14 OTUs (included representative OTU 9) in the *Arcobacteraceae* were positively correlated with migration rates (**Suppl. fig. S3a**). In a phylogenetic placement of OTU short reads, the 14 OTUs were distant from one another across the family *Arcobacteraceae* in the reference tree, although some OTUs were clustered together. For instance, OTU 8 and OTU 1074 grouped in a cluster (**Suppl. fig. S3b**). Three OTUs (OTU 27, OTU 192, and OTU 952) belonged to the genus *Malarcobacter*, one OTU (OTU 144) belonged to the genus *Halarcobacter*, and other OTUs were classified in an uncultured group (**Suppl. fig. S3b**). Although other families (see more details in **Suppl. table S3** and **Appendix**) showed no significant correlation, such as cyanobacteria belonging to the *Desertifilaceae*, they tended to increase positively along with the migration rates (**Suppl. table S3**).

**Fig. 3.**
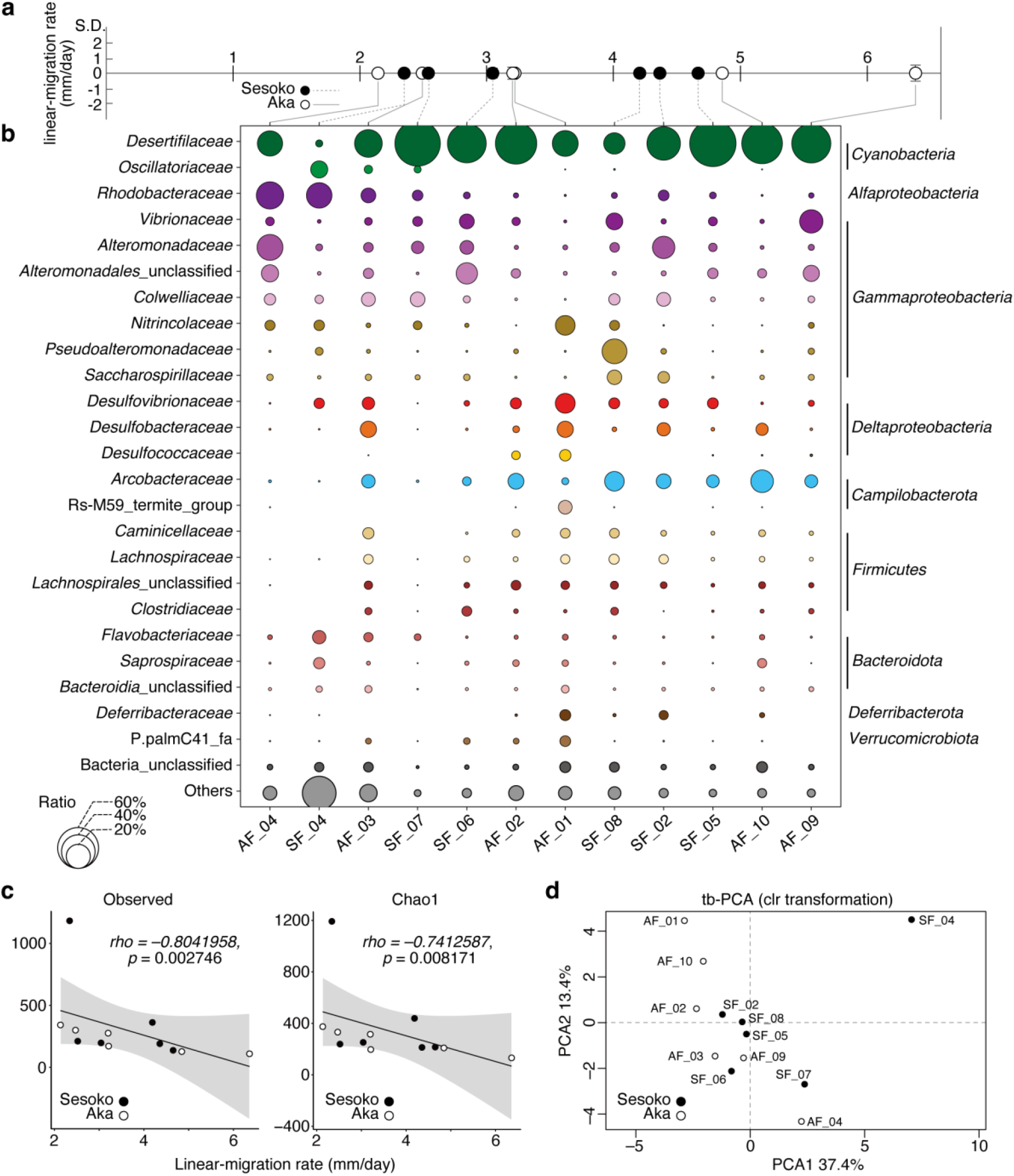
Relative abundance of bacterial communities, alpha-, and beta-diversities in BBD microbial mats showing different migration rates. Along various migration rates of BBD from two locations (**a**), bubble plots showing the relative bacterial abundance (depicted by size) of the top 24 families, unclassified bacteria and others (that average <0.5% of that relative abundance were pooled) (**b**). Significant correlations between migration rates and alpha diversities (OTU richness and Chao1 index) from BBD calculated by Spearman’s rank correlation (**c**). Grey shading shows 95% confidence intervals for the Spearman rank correlation (**c**). Transformed based principal component analysis (tb-PCA) representing the bacterial communities (family level) in BBD (**d**). Sample IDs indicated as ‘AF’ and ‘SF’ were collected from Aka Island and Sesoko Island, respectively.

**Fig. 4.**
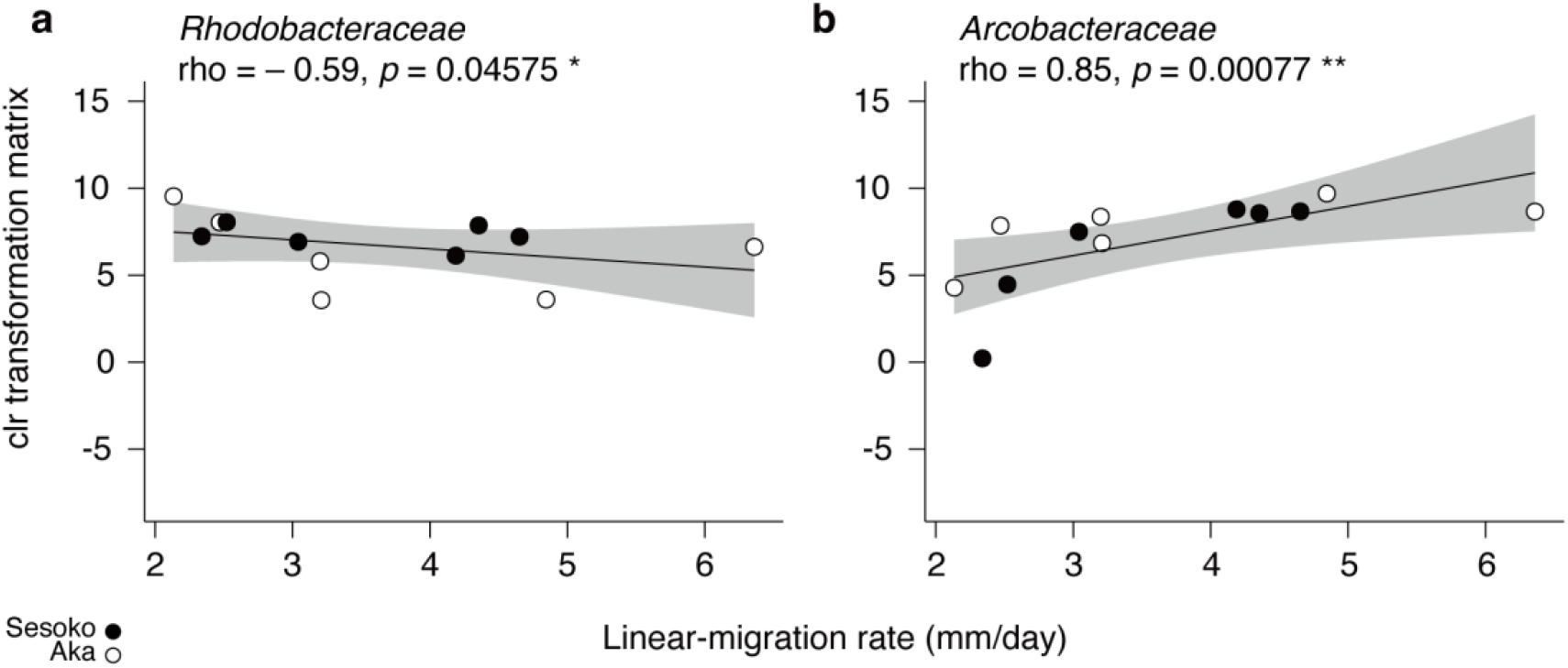
Spearman’s rank correlation with families *Rhodobacteraceae* (a) and *Arcobacteraceae* (b) to migration rates. Significant correlation marked as * *p* < 0.05 and ** *p* < 0.01. The shaded grey area represents the 95% confidence interval.

### Correlation between BBD migrations and the bacterial localizations

To study bacterial localization in BBD across distinctive migration rates, we applied FISH to inspect the bacterial distribution in the same BBD samples that were used in the bacterial community analysis. Given that sequences affiliated with the family *Arcobacteraceae* showed significant increases with the migration rate (**Fig. 4b**), we used the probe Arc94^28^, along with the broad-range bacterial probe EUB338mix^29^, to target *Arcobacteraceae*. Before conducting the FISH, we evaluated the newly designed probe that could cover the 14 OTUs in *Arcobacteraceae* using an *in-silico* analysis. Ten of the 14 OTUs, include two representatives OTUs (OTU 8 and OTU 9), were ideally matched to the specific probe Arc94 based on the phylogenetic placement analysis (**Suppl. fig. S3b**).

To visualize intact bacterial distribution and their locality in the corals, including their tissue and skeleton, we used a recently established method in which FISH^30^ was combined with undecalcified coral sectioning^31^. Although a non-specific binding signals from the FISH probe was detected in coral skeleton region (**Suppl. fig. S4**), we successfully visualized the bacterial locality within the coral tissue and intact skeleton structures (**Suppl. fig. S5**).

Many EUB338mix probe signals indicated bacterial localization in the microbial mat from the healthy coral tissue boundary to the black band. Meanwhile, EUB338mix probe signals were absent or few within the healthy tissue. The bacterial localization showed a vertically stratified structure in the cyanobacterium-dominated mat where cyanobacteria covered the uppermost layer, while many bacteria spread around necrotic coral tissues and symbiotic dinoflagellates below the cyanobacterial layer (**Fig. 5a**). In serial sections, assemblages and individual cells of *Arcobacteraceae* were also observed under the cyanobacterial layer using the specific probe Arc94 (**Fig. 5b**). While bacterial assemblages from EUB338mix signals showed a variety of individual shapes (**Fig. 5c**), *Arcobacteraceae* assemblages showed an almost same cell morphology (rod-shaped bacteria, **Fig. 5d**). Moreover, the distribution pattern of *Arcobacteraceae* indicated their close associations with coral necrotic tissues (**Fig. 5d**).

**Fig. 5.**
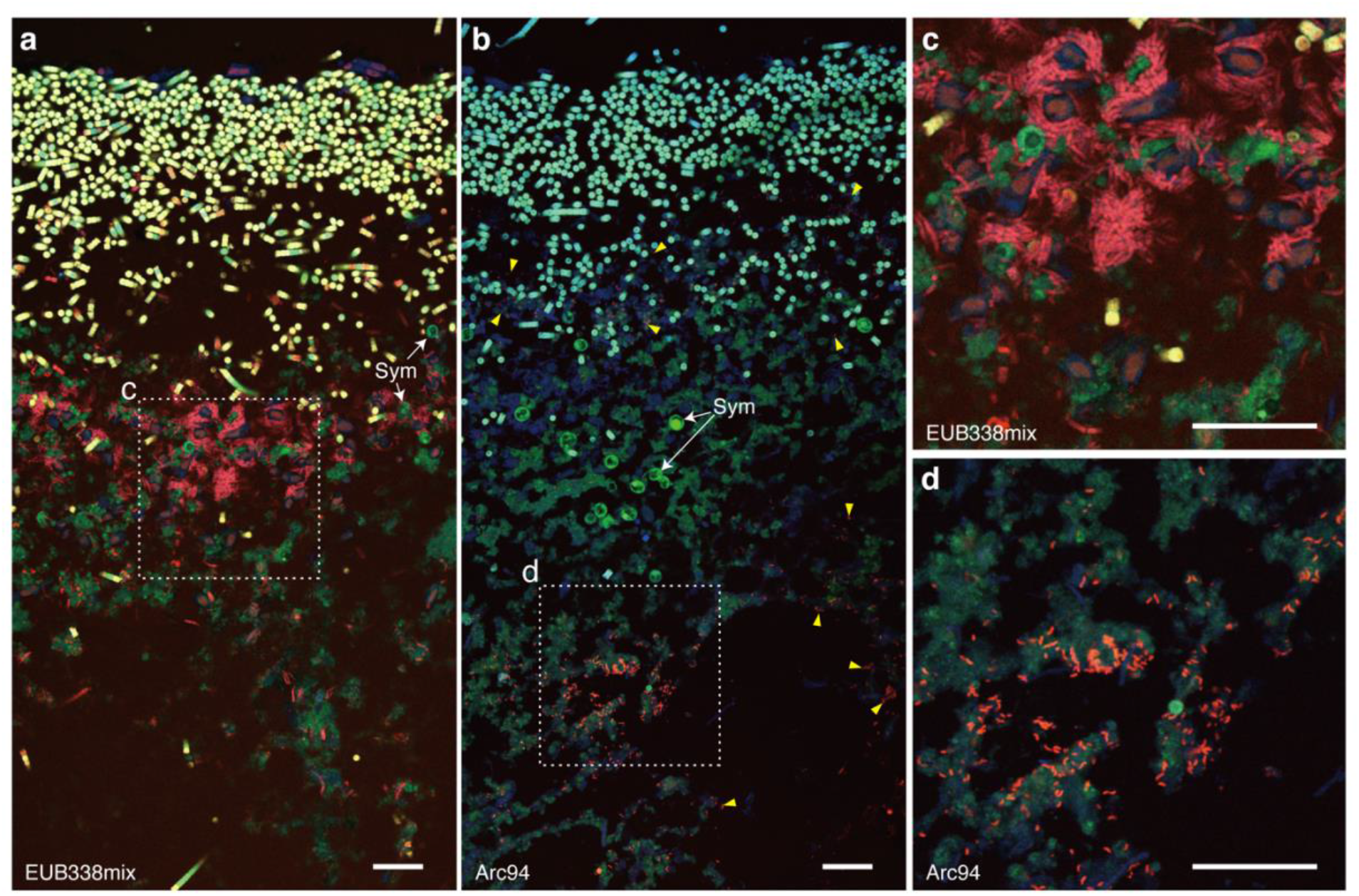
Fluorescence *in situ* hybridization (FISH) using EUB338mix and Arc94 probes labeled with Cy3 showing respective bacterial localizations of all bacteria (a and c) and *Arcobacteraceae* (b and d) in serial cross sections of BBD specimens. Confocal merge micrographs of the microbial mat structure of BBD showing autofluorescence (blue indicates coral tissue and green indicate chlorophyll from symbiotic dinoflagellates [Sym] and cyanobacteria) and the hybridized bacteria with probe labeling Cy3 (red) (**a-d**). Cyanobacteria is shown in a yellow color that merged two channels from autofluorescence of chlorophyll and bacterial signals with Cy3 (**a** and **c**). Dotted lines in **a** and **b** delineate magnified, close-up regions (**c** and **d**). Hybridized signals of *Arcobacteraceae* shows their assemblages and dispersed cells (dotted line and yellow arrowed heads in **b**). Scale bars represent 30 µm.

To examine the relationship between bacterial locality and migration rate, the areas of bacterial presence of all bacteria and *Arcobacteraceae* were spatially quantified in the upper, middle, and bottom layers of the microbial mats (each vertical interval 1 mm, **Fig. 6a**) and compared against the migration rates. Given the SEM observations and bacterial community analysis showing that the dominant cyanobacteria always occurred in the upper layer, the area of filamentous cyanobacteria was excluded in this quantitative analysis (hereafter the “area of all bacteria” is defined as having a distribution of bacteria, excluding cyanobacteria). FISH was performed using the EUB338mix and Arc94 probes on serial sections so that the distributions of target bacteria could be optimally compared (**Fig. 6b** and **c**). The detection area of all bacteria, which was measured with the EUB338mix probe was larger than the area of *Arcobacteraceae*, which was measured with specific Arc94 probe (**Fig. 6d**). Both FISH signals were detected in all three layers across different migration rates (**Fig. 6e-j**). In the surface layer, detection of all bacteria and *Arcobacteraceae* did not show any correlations with the migration rate (**Fig. 6e** and **f**). Only in the middle layer did the area of *Arcobacteraceae* have a positive correlation with the migration rate (adj. R-squared 0.04 and *p* value < 0.05) (**Fig. 6g** and **h**). In contrast, all bacteria showed a significantly positive correlation between the area and the migration rate only in the bottom layer (adj. R-squared 0.06 and *p* value < 0.05) (**Fig. 6i** and **g**).

**Fig. 6.**
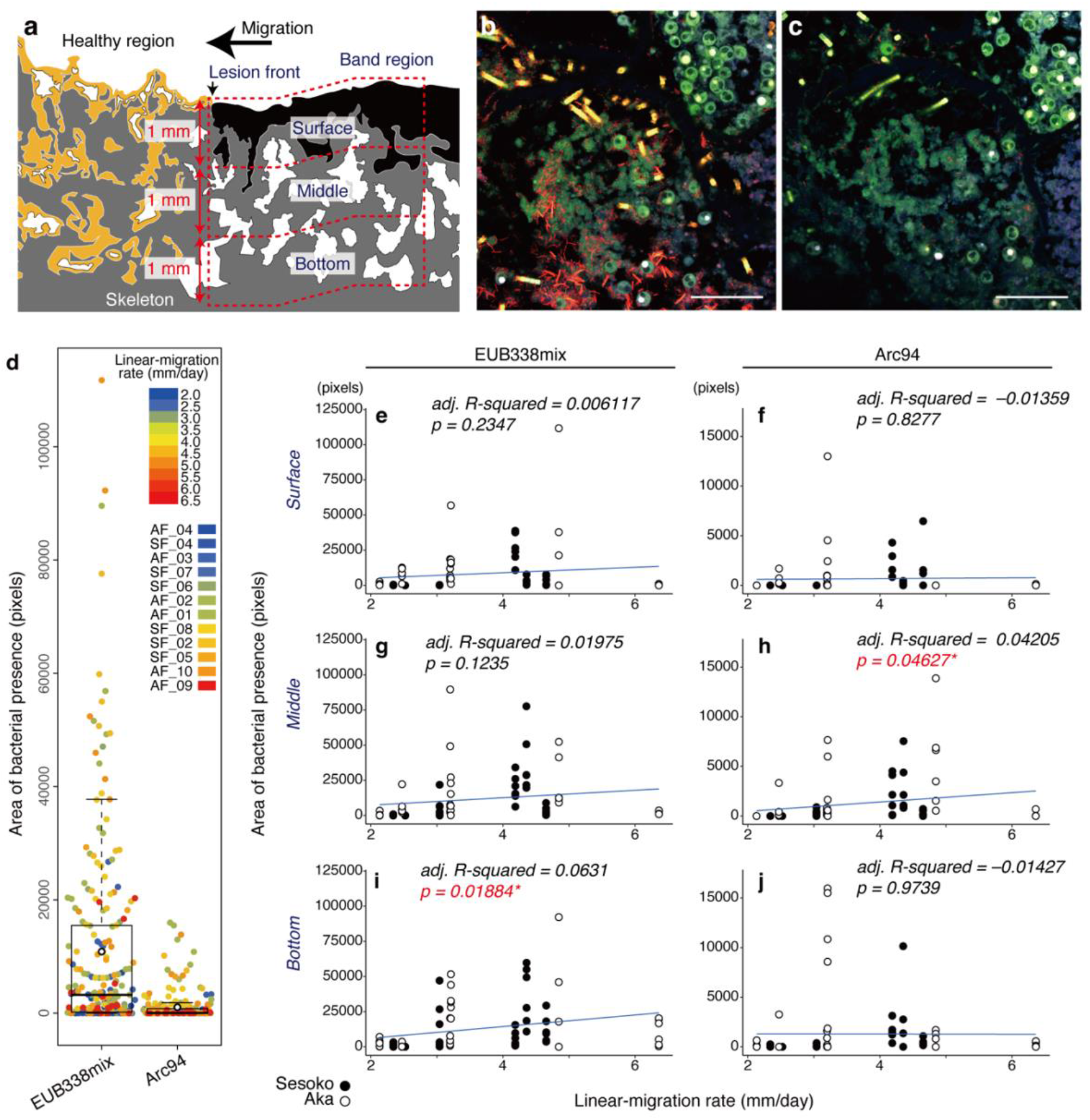
Quantitative area analysis of the spatial bacterial localizations of all bacteria (except filamentous cyanobacteria) and *Arcobacteraceae* in BBD mats. Schematic illustration showing three layers (surface, middle, and bottom; each vertical interval is 1 mm) in BBD mats for the spatial analysis (**a**). Confocal merge micrographs showing spatial bacterial localization (red) of all bacteria (**b**) and *Arcobacteraceae* (**c**) in two serial sections; hybridized with EUB338mix and Arc94 probes, respectively. Scale bars indicate 50 µm (**b** and **c**). Dot plots merged with box plot showing the entire area of both all bacteria (except filamentous cyanobacteria) and *Arcobacteraceae* detections in the various migration rates (depicted by color;the sample IDs ‘AF’ and ‘SF’ indicatecollection from Aka Island and Sesoko Island, respectively) (**d**). Linear regressions showing the associations between migration and areas of bacterial presence (all bacteria [**e, g**, and **i**] and *Arcobacteraceae* [**f, h**, and **j**]) in the surface (**e** and **f**), middle (**g** and **h**), and bottom layers (**I** and **j**). Significant correlation is shown in red, with * *p* value < 0.05.

## Discussion

In this study, we successfully demonstrated that the migration (i.e., a proxy for virulence) of cyanobacterial-dominated microbial mat is linked to both bacterial composition and spatial localization using a novel combination of 16S bacterial profiling, undecalcified sectioning, and FISH. The BBD bacterial community in this study, which showed a unique migration pattern as a band-like biofilm, was composed of groups that became less diverse as the migration rate increased. In the bacterial community analysis, we also found that the relative abundances of the families *Rhodobacteraceae* and *Arcobacteraceae* had negative and positive correlations with the BBD migration rates, respectively. Moreover, as the BBD-migration rate increased, the population of *Arcobacteraceae* significantly increased in the middle layer, while all non-cyanobacterial populations increased in the bottom layer.

### The involvement of the Arcobacteraceae in the BBD-virulence

Our results suggest that *Arcobacteraceae* (= *Arcobacter sensu lato*^32–34^) are profoundly implicated in the pathogenicity of BBD. The presence of *Arcobacter* in BBD has been commonly reported in a wide array of geographical regions in the Caribbean, the Indo-Pacific and the Red Sea^27,35–37^. *Arcobacteraceae* are aerotolerant, psychrotrophic^34^, and ubiquitous in environments and animals^38^,aerotolerant, and psychrotrophic^34^. Considering the spatially (especially vertically) and temporarily heterogeneous oxic condition within the BBD mat^39^, our finding that *Arcobacteraceae* populations appeared in all layers of BBD with low to high migration rates suggests that the *Arcobacteraceae* associated with BBD are aerotolerant.

A member of *Arcobacteraceae* isolated from marine environments has been identified as either an autotrophic or chemolithoheterotrophic sulfide oxidizer through *in vitro* and *in silico* analyses^40–42^. In previous BBD study, the relative abundance of *Rhodobacteraceae* decreased during the BBD-development transition from the BBD-precursor, called a ‘cyanobacterial patch’ (CP), which progressed at a slower rate, whereas the relative abundance of *Arcobacter* spp. increased as BBD developed with increased virulence^19^. The contribution of *Rhodobacterales* in BBD has been confirmed by profiling using a key functional gene (*soxB*) for sulfide oxidation in BBD. However, the relative abundance of *soxB* genes affiliated with *Rhodobacterales* increased more in the BBD mat compared to CP^16^. *Arcobacter*-derived *soxB* genes were not detected, presumably because the universal primers for *soxB* genes are incapable of identifying sequences from the genus *Arcobacter*^43^. In fact, a shotgun metagenome sequencing study, free from PCR biases, reported *Arcobacter* as major members of SOB in BBD^36^. Taken together, there is a strong possibility that *Arcobacter* plays a role in the sulfide oxidization stage of the sulfur cycle in the BBD consortium. In addition, since sulfur oxidizing *Arcobacter* is highly resistant to high hydrogen sulfide and low oxygen levels and thus effectively competes with other co-occurring SOBs^44^, our results indicate that a functional niche of the sulfide oxidization in the BBD may have transferred from *Rhodobacteraceae* to *Arcobacteraceae* as BBD-virulence became higher.

Our results from FISH showed that the population of *Arcobacteraceae*, particularly in the middle layer of the BBD mat, positively correlated with the migration rates. Although the vertical chemical distributions of hydrogen sulfide and oxygen vary dynamically in the BBD mat from daytime (low hydrogen sulfide and oxygen vertically decreases with mat depth) to nighttime (hydrogen sulfide vertically increases and the mat becomes entirely anoxic)^39^, *Arcobacter* could be capable of mediating within any condition and maintaining their abundance^45^. Furthermore, *Arcobacter* is a highly motile microaerophile, which appears to be typical of organisms living at oxic-anoxic interfaces^44^. Considering the chemical condition in BBD at the daytime^39^ that we collected samples, the middle layer of BBD mat could allow the *Arcobacteraceae* to proliferate and adapt to the environmental conditions, including hydrogen sulfide and oxygen levels, and simultaneously provide the sulfate for SRB by the sulfide oxidization and play a role in promoting or changing the consortium’s migration that could associate with the BBD-virulence. Furthermore, given that the genus *Arcobacter* also has the complete set of genes responsible for assimilatory sulfate reduction according to the whole-genome prediction^45^, future studies are required to confirm the role of *Arcobacteraceae* in not only sulfide oxidation but sulfate reduction in the BBD mat.

*Arcobacteraceae* in BBD could also behave as pathogens that directly impair coral tissues. *Arcobacter* has also been abundantly detected in diseased lesions of many coral diseases other than BBD, such as White Pox^46^, White Syndrome^47^, Brown Band Disease^47^, and Stony Coral Tissue Loss Disease^48^ across the world’s coral reefs. *In vitro* human and animal cell culture assays have shown that *Arcobacter* have significant virulence when it comes to colony formation and the establishment of infection in host tissues^49,50^, and it is also known that most *Arcobacter* are equipped with virulence genes such as *ciaB, irgA*, and *cadF*^51,52^. Therefore, the presence of pathogenetic *Arcobacter* in marine organisms warrants further focused research.

### Bacterial dynamics in BBD with various migration rates

Biofilms have been involved in many clinical infections, and accumulated evidence shows that biofilms contribute to pathogenesis, especially chronic infection^53^ and oral plaque^54^. Of the pathogenic biofilms, the most well-studied systems include human oral plaque. After a partial clearance of the polymicrobial biofilm by physical removal, the colonization cycle repeats itself in the same general spatiotemporal successional transition until a mature community of microbes is repopulated^55^. During the transition, early bacterial colonization and composition vary among individuals, but the most abundant genera are usually conserved. Abrupt or subtle changes in bacterial composition are often thought to promote disease-associated phenotypes^56^. Specific polymicrobial associations in human periodontitis may exacerbate disease severity and progression^56^. In contrast, the variation of BBD migrations could not be linked to the certain bacterial communities that appeared in BBD with a high migration rate, although the alpha diversity was negatively correlated. Indeed, the bacterial family *Vibrionaceae*, which includes some species known to be coral pathogens^57,58^, appeared regardless of the migration gradient (**Fig. 2b**). Nevertheless, a trend of decreasing *Rhodobacteraceae* and increasing *Arcobacteraceae* among BBD with a high migration rate was certainly confirmed.

In the FISH experiment, bacteria in the BBD microbial mat showed spatial distribution patterns with various migration rates. Besides the population of *Arcobacteraceae*, all bacterial populations in the bottom layer positively correlated with the BBD migration rate. Combined with the bacterial community results, our microbial visualization suggested that the bacterial community structure linked to high BBD-virulence thrives under strong selection that excludes other bacteria, especially in the bottom layer. In the bottom layer of BBD, where low oxygen or anoxic conditions are anticipated, even in the daytime^14^, a niche is likely provided for both SRB and anaerobic heterotrophs.

This study demonstrates for the first time the correlation between changes in bacterial composition/spatial localization and the migration rate of BBD. In short, this study provides new insights into the microbial dynamics of BBD and indicates that the microbe-mediated pathogenesis model in BBD is more complicated than previously thought. However, our results strongly indicate that *Arcobacteraceae* could be one of the key foundations in the sophisticated bacterial community structure of BBD. Given the positive correlation between *Arcobacteraceae* and BBD-virulence, we propose *Arcobacteraceae* to be a potential biomarker for BBD-virulence.

## Methods

### Study site and sample collections

All samples of BBD-affected encrusting *Montipora* colonies were collected from reefs around Sesoko Island (26°38’35.2”N, 127°51’49.5”E) and Aka Island (26°12’00.0”N, 127°16’45.0”E) in Okinawa, Japan (**Suppl. fig. S1**) in 2014 and 2015. The sampling sites are located approximately 70 km away from each other and have open-ocean between them.

For scanning electron microscope (SEM) observation, 18 BBD samples (n = 9 from each location, approximately 1cm square) were obtained from each location in August 2014, immediately fixed by 2% glutaraldehyde (GA) in 10 mM phosphate buffered saline (PBS, pH 7.4) on ice, and sequentially stored at 4°C.

For bacterial community analysis and FISH, 12 BBD samples (n = 6 samples from each location) were collected in August 2015 and cut into two specimens each. One specimen was punched from the black band by sterilized leather punch (4 mm circle diameter), and the band was stored in 300µl of 100% ethanol at −80°C for bacterial community analysis. The second specimen (approximately a 2 cm square) was immediately fixed in 4% paraformaldehyde (PFA) in PBS (Wako, Japan) at 4°C for 8 hours, rinsed with 70% ethanol three times, and stored in 70% ethanol at 4°C for FISH below.

To determine the linear-migration rate of BBD on each colony before collection, photographs were taken of the front of the BBD-lesion on a flat region of the colony, both three-days before the sampling and on the day of the sampling. Average linear-migration rates (mm/day) of BBD were calculated at five random points on the region using Fiji software^59^. All samples were collected from the region where the linear-migration rates were measured. We also measured depths for each BBD-infected colony collected.

### Scanning electron microscope (SEM)

The GA-fixed samples (n = 18) were gently rinsed in PBS for 15 min three times, and dehydrated in an ethanol/water gradient series (20, 40, 60, 70, 90, and 100% once, and abs. ethanol three times) for 20 min each at room temperature. Dehydrated GA-fixed samples were immersed in abs. ethanol/t-butyl alcohol graded series (7:3 and 1:1 for 15 min each), transferred to t-butyl alcohol at above 25.5°C for 60 min, and placed in a refrigerator after changing out old t-butyl alcohol with new alcohol. The samples were dried using a freeze-drying devise (VFD-21S, SHINKKU VD, Japan) for 3 hours, and coated with a platinum/palladium alloy in ion-sputter (E-1010, HITACHI, Japan). Observation and image acquisition were conducted by a scanning electron microscopy (SEMS-3500N, HITACHI, Japan). We observed the microorganisms in the surface area within 1 mm from the border with healthy coral tissue to the BBD mats, and determined the presence or absence of microorganisms by randomly photographing. A total of 42 images (6 images at 100x magnification, 18 images at 1000x magnification, and 18 images at 2000x magnification) were taken from each BBD lesion.

### Bacterial community analysis

The BBD-punched samples (n = 12) in 300 µl of 100% ethanol, were directly added 120 µl of nucleotide-free water and 12 µl of 3M sodium acetate, mixed by vortexing, and stored at −20°C for 45 min. After a centrifugation at 18,000 x *g* for 20 min, the pellet was resuspended in 300 µl of prechilled 70% ethanol, centrifuged again by same condition above, removed from the solution, and dried. Genomic DNA was extracted from the pellet by DNeasy Blood & Tissue kit (Qiagen), which included the addition of a lysozyme-based enzymatic lysis step (buffer: 20 mg/ml lysozyme, 20 mM Tris-HCl [pH 8.0], 2 mM EDTA [pH 8.0], and 1.2% Triton X-100) according to the manufacturer’s protocol.

The genomic DNA was used as a template for amplification of the V4 region of bacterial 16S rRNA gene using the universal primer set (515F and 806R, **Suppl. table S4**) and the following PCR procedure: 35 cycles of 30 sec at 95ºC (denaturation), 30 sec at 51ºC (annealing), 30 sec at 72 ºC (extension), followed by an additional extension for 5 min at 72ºC. Then, index sequences were added by PCR using Nextera XT index kit (Illumina, US). The amplicons were sent for sequencing with a paired-end read chemistry by a MiSeq sequencer (Illumina, US). The sequencing reads are summarized in **Suppl. table S5**.

The paired-end reads from each sample were merged in the software QiimeI using MacQIIME v.1.91^60^. The obtained merged sequences were filtered in the software MOTHUR v.1.39.5^61^ using the following criteria: 1) read lengths between 249 and 256 bp; 2) read quality score average 27; and 3) homopolymer read length < 8 bp. We further detected chimeric sequences using the USEARCH algorithm v.11.0.667^62^ and subsequently excluded chimeric sequences. We implemented USEARCH with a 97% similarity cut-off to cluster OTUs. The OTUs were assigned to known taxonomic groups by mapping onto the Silva SSU r138.1 database^63^ using the MOTHUR with a cut-off value of 80. After removing sequences from Eukaryota, Archaea, unknown and chloroplast, a total of 1,593 bacterial OTUs were assigned to 255 families (1401 OTUs), unclassified bacteria (151 OTUs), uncultured bacteria (31 OTUs in 10 orders), uncultured families (four OTUs in three classes), and unknown families (six OTUs in two orders).

Statistical analyses for the bacterial community data set were performed using the statistics software R v4.0.2^64^, incorporating the R package phyloseq v.1.34.0^65^ and vegan v. 2.5-7^66^. Alpha diversity indices (OTU richness and Chao1) were calculated using the function “estimate_richness” within the phyloseq, and the correlation along migration rates was using Spearman’s rank correlation in R v4.0.2^64^. For beta diversity analysis, the phyloseq data at the family level was transformed using the Centered Log Ratio (clr) transform function in a R package microbiome v. 1.12.0^67^. Transformation based principal component analysis (tb-PCA) using Euclidean distance was performed with the function “ordinate” from the R package phyloseq. To assess the effect of the migration gradient on community structure, generalized additive models with integrated smoothness estimation from the R package mgcv v.1.8-36^68^ and Mantel test from the R package vegan were applied for the fitness and correlation of the migration gradient. Subsequently, a partial correlation analysis was determined using clr matrix of each representative bacterial family and BBD linear-migration rate calculated using the Spearman’s rank correlation test in R.

### Fluorescence in situ hybridization (FISH)

According to the partial correlation analysis, family *Arcobacteraceae* included 14 OTUs that were positively correlated with the BBD migration. Therefore, we performed FISH on 12 samples using the specific probes, EUB338mix^29,69^ for most bacteria, and Arc94^28^ for *Arcobacteraceae* (**Suppl. table S4**), with a undecalcified thin section method, after estimating the probe specificity for the 14 OTUs.

### Phylogenetic placement of OTU short reads of Arcobacterease

To estimate the probe specificity for the 14 OTUs from *Arcobacteraceae*, 653 reference sequences assigned to family *Arcobacteraceae* comprised of the genera *Arcobacter, Malaciobacter, Halarcobacter, Pseudarcobacter, Poseidonibacter* and uncultured *Arcobacteraceae* were retrieved from the Silva SSU r138 database^63^. The reference sequences were also tested for matching with the Arc94 probe ^28^ for *Arcobacter* using TestProbe 3.0 (https://www.arb-silva.de/search/testprobe/) and the probe estimated a coverage of *Arcobacteraceae* at 78.0% (509 out of 653 reference sequences). For construction of the reference tree, the reference sequences were aligned using the software infernal v.1.1.3^70^ and a maximum likelihood tree was constructed in RAxML-NG v.1.0.2^71^ with model GTR+G. Then, 14 OTU sequences were aligned to the reference multiple sequence alignment using the program PaPaRa core v.2.5^72^, subsequently placed with the highest estimation score (like_weight_ratio) on the reference tree by the software EPA-ng v.0.3.6^73^, and then the tree was visualized using interactive Tree of Life (iTOL) v.4^74^ for the evaluation of the probe specificity.

### FISH on undecalcified coral thin sections

The undecalcified section attached to adhesive film performed three serial sections from the PFA-fixed samples for coral specimens, as described previously^31^, at 5 µm in thickness with a tungsten carbide blade (SL-T30, SECTION LAB Co. Ltd) using a cryostat system (Leica CM1850). The undecalcified sections attached to the adhesive film were immerged in 100% ethanol for 30 sec to remove the compound and air-dried. FISH was carried out as described previously^30^ with slight modifications including a non-dewaxing step and a non-immerging step with HCl solution. Each serial section was subjected to FISH using three probes that were labeled with the Cy3 fluorochrome: 1) EUB338mix^29,69^,2) Non338^75^ at a hybridization buffer containing 30% formamide, and 3) Arc94^28^ at that 20% formamide (**Suppl. table S4**). The hybridized sections attached to adhesive film were air-dried, collected on a glass slide (S2441, MATSUNAMI), and then a coverslip was mounted on the slide in an antifade solution (Fluoromount/Plus, Diagnostic BioSystems).

Image acquisition was conducted with a 40x magnification objective lens (HC PL APO 40x/1.30 OIL CS2) in a confocal microscope (TCS SP8, Leica). Cy3 fluorochrome and chlorophyll (for Symbiodiniaceae and cyanobacteria) were excited at 552 nm (2.0%) and detected in an emission range of 571–582 nm with HyD (standard mode and gain 100) and of 650–696 nm with PMT (gain 700). Coral autofluorescence was also excited at 405 nm (1.5%) and detected in an emission range of 460–510 nm with HyD (standard mode and gain 161). In order to quantitatively analyze the area for the presence of bacteria, six images (each size 1024 × 1024 pixels) were acquired from each of the three regions of the BBD cross section: upper, middle, and bottom layers (each vertical interval 1 mm, **Fig. 6a**). Fluorescence signals bounded by each probe (EUB338mix and Arc94) were identified with whither specific signals or non-specific probe bindings (using Non338 probe), according to the criteria as detailed by Wada et al^30^.

Image processing was performed using the software Fiji^59^ after exporting the images of Cy3 fluorochrome and chlorophyll to 8-bit greyscale TIFF images using the software LAS X (Leica). Each greyscale image was converted to a binary image by a thresholding function using the algorithm “Max entropy”^76^. To analyze the presence of bacteria other than cyanobacteria in the Cy3 signal of EUB338mix probe, the region of cyanobacteria in the chlorophyll signal was subtracted from the binary image of the EUB338mix signal by the function “subtract” and manual curation. The binary image was filtered to reduce the noise by the function “Remove Outliers” (Radius 1.5 and threshold 50). In addition, since the signals from Cy3 fluorochrome channel also contained non-specific fluorescence from the skeleton regions (**Suppl. fig. S4**), the fluorescence from skeleton regions was manually removed from the binary image. Based on the binary images, areas of total bacteria, except for cyanobacteria and *Arcobacteraceae*, were calculated in pixels. Each area from three layers was tested to measure the correlation with BBD linear-migration rate using Spearman’s rank correlation test in the statistics software R v4.0.2^64^.

## Data availability

All sequencing data produced in this study have been submitted to the DDBJ Read Archive under accession number DRA010783.

## Acknowledgement

This research was supported by Grant-in-Aid for JSPS Fellows DC1 26•6085, and funding from Academia Sinica. N.W. kindly thanks Dr. Kataaki Okubo, Dr. Yukika Kawabata-Sakata and the late Mr. Kiyoshi Nakasone for the technical support in confocal microscope imaging and encouraging N.W.’s motivation for his PhD. N.W. also thanks both the late his dad and his mom for opening doors of opportunities for him to research. N.W. sincerely hope their journey are going well in heaven.

## Contribution

N.W. and N.M. conceived the study. N.W., Y.U., N.A., K.T., S.H., and S.K. conducted the filed sampling including the measurement of BBD-migration rate. N.W. and Y.U. performed the SEM observation. N.W., A.I. and Y.Y. conducted molecular processing, and bioinformatic and statistics analysis on sequence data. N.W. carried out the histological work, FISH observation and statistics analysis. N.W. and S.-L. T. had major contribution in the manuscript writing and the figure making. A.I., Y.S. and N.M. contributed to writing and editing the manuscript. All authors critically reviewed.

## Conflict of interest

The authors declare they have no conflict of interest.

## Notes

### Competing Interest Statement

The authors have declared no competing interest.

