## Supplementary figures for "Microbial mat compositions and localization patterns explain the virulence of black band disease in corals"

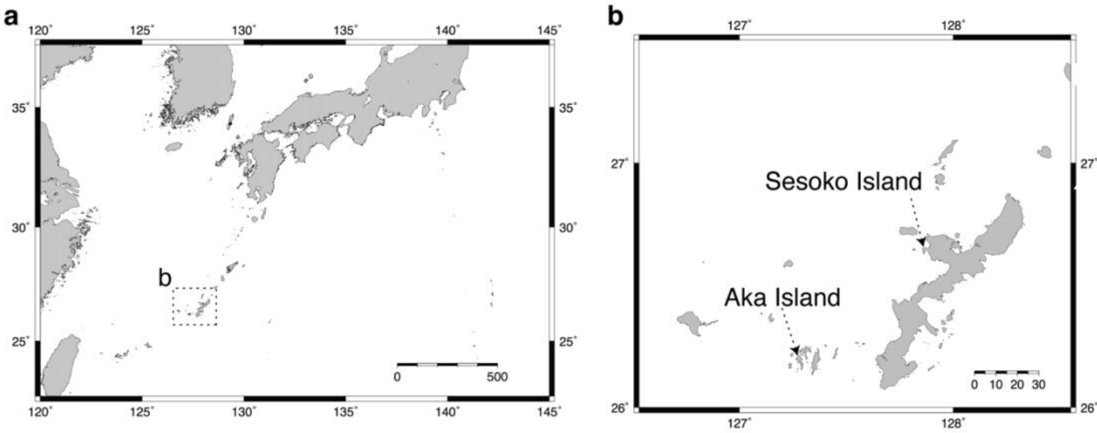


**Suppl. fig. S1 Map showing the locations of two study sites in Okinawa, Japan (a-b).** Sesoko Island and Aka Island are separated by more than 70 km (**b**).

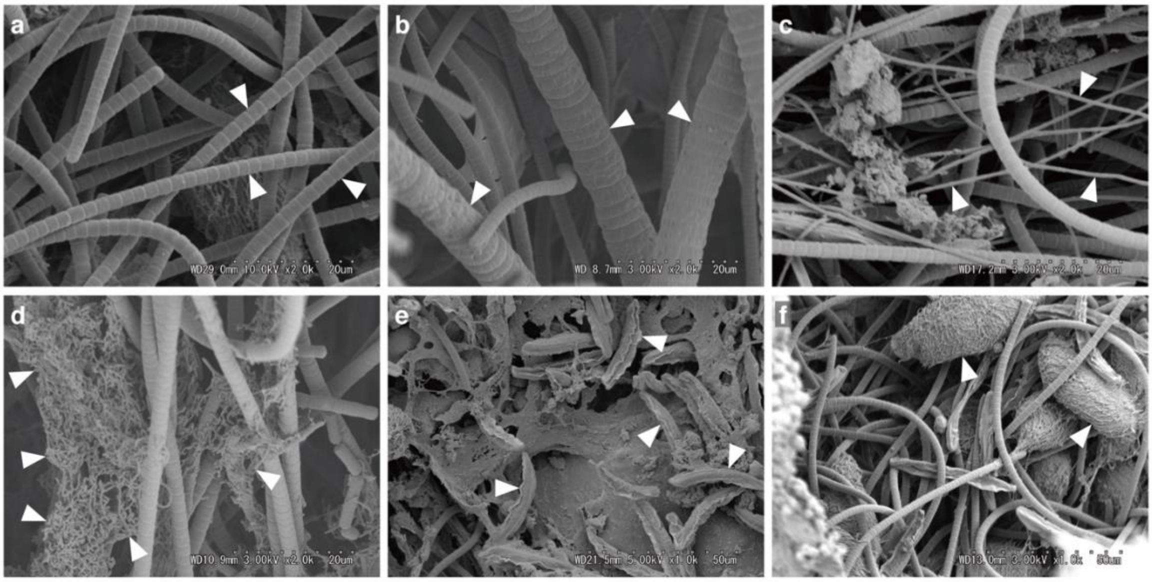


**Suppl. fig. S2 Representative microbes on the BBD surface.** SEM images of **a**) filamentous cyanobacteria (cell length: 2.69 ± 0.04 µm long and 2.11 ± 0.32 µm broad), **b**) thick cyanobacteria (2.63±0.16 µm long and 8.56±0.11 µm broad), **c**) filamentous microorganisms (0.76±0.32 µm broad**)**, **d**) bacterial aggregation, and **e-f**) two kinds of ciliates (Type A(**e**): 31.3±1.07 µm long and 5.53±0.18 µm broad, and Type B (**f**): 40.8±0.91 µm long and 17.5±0.51 µm broad). The scale bars indicate 20 µm (**a**-**d**) and 50 µm (**e**-**f**).

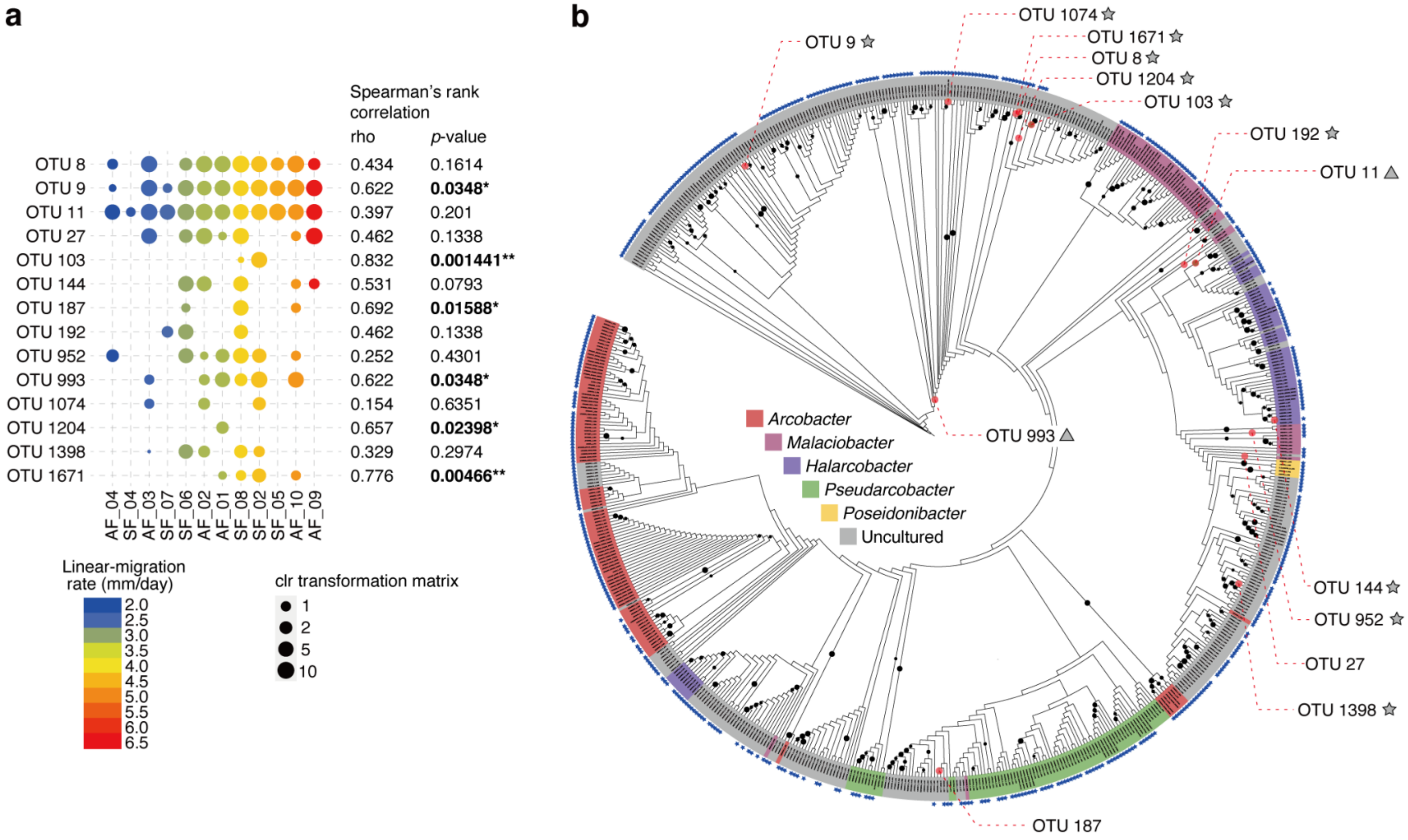
a
Spearman's rank correlation
rho
p-value
OTU 8 0.434 0.1614
OTU 9 0.622 0.0348*
OTU 11 0.397 0.201
OTU 27 0.462 0.1338
OTU 103 0.832 0.001441**
OTU 144 0.531 0.0793
OTU 187 0.692 0.01588*
OTU 192 0.462 0.1338
OTU 952 0.252 0.4301
OTU 993 0.622 0.0348*
OTU 1074 0.154 0.6351
OTU 1204 0.657 0.02398*
OTU 1398 0.329 0.2974
OTU 1671 0.776 0.00466**
AF_04
SF_04
AF_03
SF_07
SF_06
AF_02
AF_01
SF_08
SF_02
SF_05
AF_10
AF_09
Linear-migration rate (mm/day)
2.0
2.5
3.0
3.5
4.0
4.5
5.0
5.5
6.0
6.5
clr transformation matrix
1
2
5
10
b
OTU 9
OTU 1074
OTU 1671
OTU 8
OTU 1204
OTU 103
OTU 192
OTU 11
OTU 993
Arcobacter
Malaciobacter
Halarcobacter
Pseudarcobacter
Poseidonibacter
Uncultured
OTU 144
OTU 952
OTU 27
OTU 1398
OTU 187

**Suppl. fig. S3 Partial correlation between clr transformation matrixes of each OTU in *Arcobacteraceae* and BBD-progressions (a) and phylogenetic placement of 14 OTU sequences on reference tree of family *Arcobacteraceae* (b).** (**a**)The clr transformation matrixes of each OTU were calculated spearman's rank correlation with liner-progression rates and showed significant that are marked in bold with * $p < 0.05$ and ** $p < 0.01$ (**a**). The bubble chart showing clr transformation matrix (depicting by size) and progression rates (depicting by color). (**b**) The reference tree was generated from an alignment of 16S rRNA genes from a total of 653 sequences, which comprise five genera and uncultured group that assigned by Silva SSU ref v138 database, using infernal (**b**). The tree was constructed with Maximum Likelihood method using RaXML_NG with 200 bootstrap replicates (only bootstrap values greater than 50% are denoted as black cycle at the node). For the outmost small bule star marks, it indicates that specific probe Arc94 can match to the corresponded sequences (matched 509 sequences) that were calculated by Silva TestProbe. The phylogenetic placements of OTU sequence were estimated by EPA-ng on the reference tree and defined with high score of like_weight_ratio, when the argonium calculate multiple places on the reference tree (the score defined in the ranging from 0.15 to 0.99). The OTUs labels with a grey star and a grey triangle on the tree indicate probe-match (a single node corresponding to a sequence that matched the probe is defined) and unclear (contains multiple nodes including miss-matched probe), respectively. The OTUs labeled without a symbol indicate that a single node is defined but miss-matched with the probe.

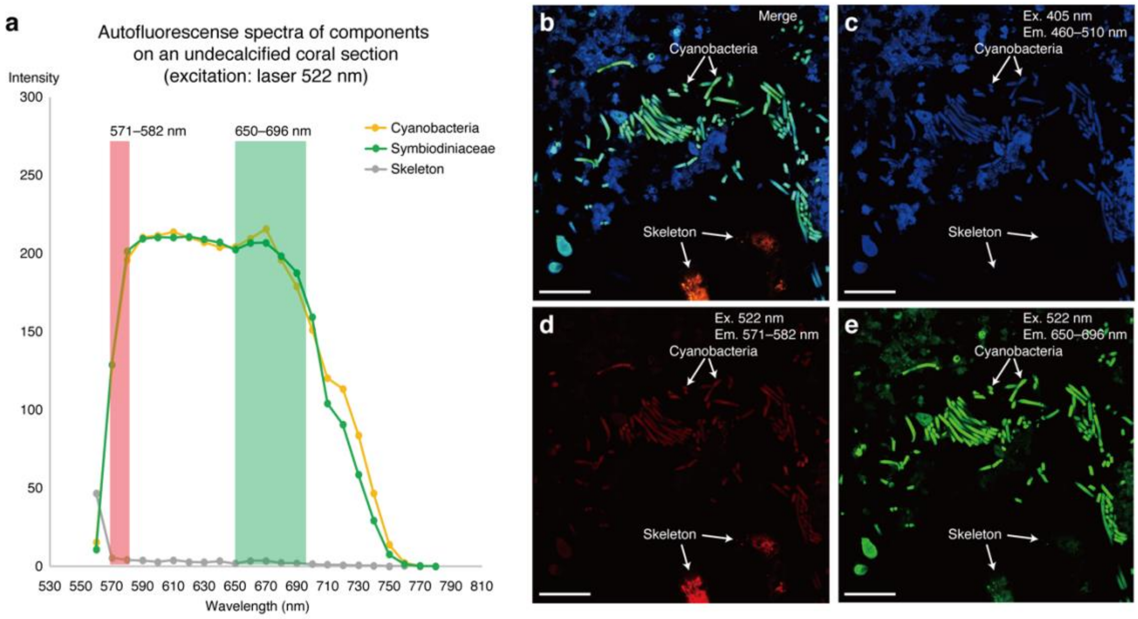


**Suppl. fig. S4 Autofluorescence spectra at excitation of a laser 552 nm on an undecalcified coral section (a) and fluorescence *in situ* hybridization image using a negative control probe Non338 (b).** Autofluorescence spectra (**a**) of cyanobacteria, Symbiodiniaceae and skeleton on a FISH-untreated undecalcified coral section, excited by a laser 522 nm (5% intensity), using a Lambda scan function of a confocal (TCS SP8, Leica). FISH images using the probe Non338 labeled with Cy3 showing a merged image (**b**) from three different channels (**c**-**e**). Given that skeleton is not shown autofluorescence, the FISH result indicating the non-specific binding on skeleton region from Non338 probe. Scale bars indicate 50 µm.

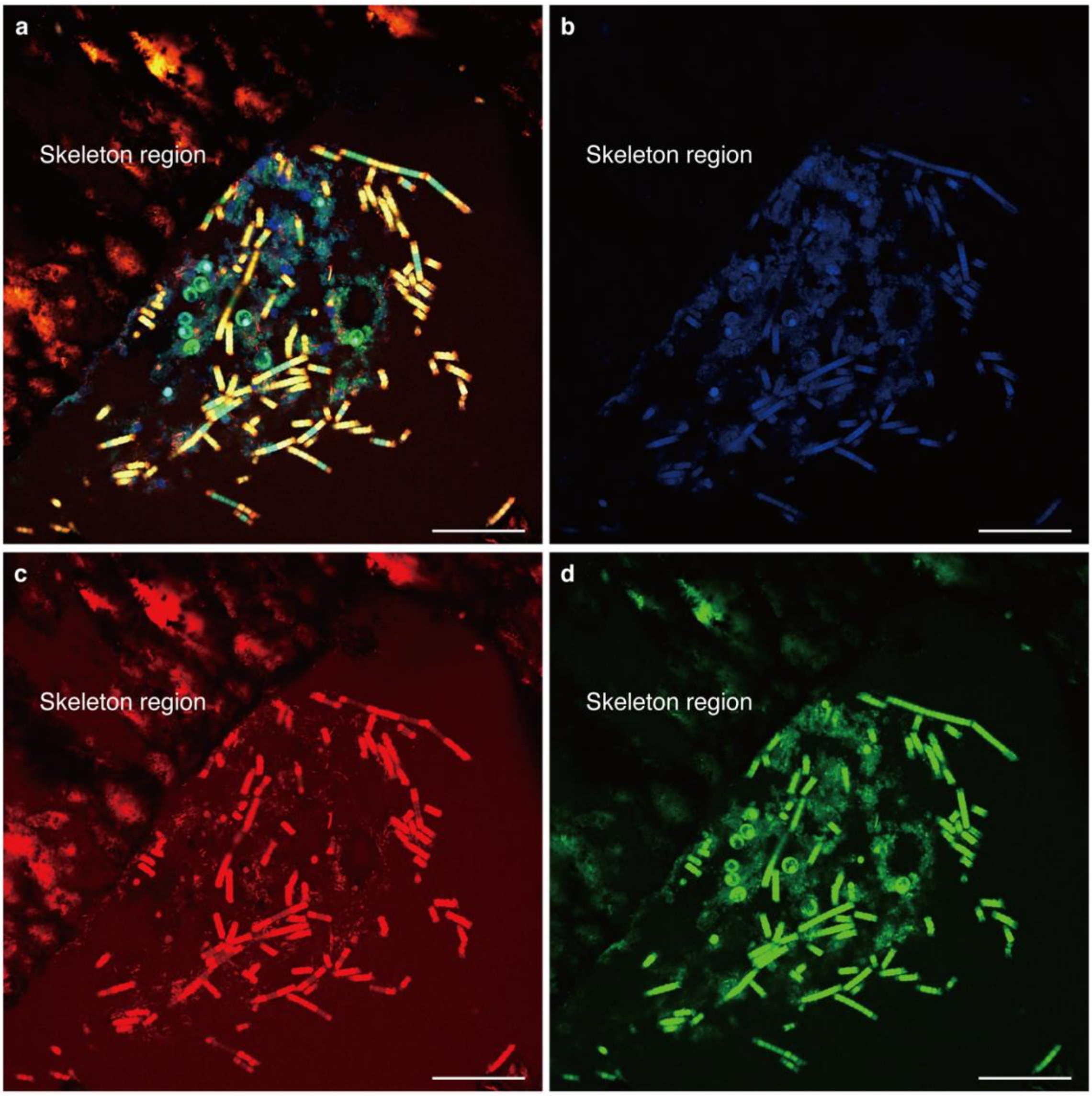


**Suppl. fig. S5 Fluorescence *in situ* hybridization images showing EUB338mix probe (labeled with Cy3) binding with bacteria and cyanobacteria on a undecalcified coral section.** Merged image (a) is displayed signals from mainly coral autofluorescence (**b**, blue) excited at leaser 405 nm (emission: 460–510 nm), Cy3 fluorochrome (**c**, red) excited at 552 nm (emission: 571–582 nm), and mainly chlorophyl (**d**, green) excited at 552 nm (emission: 650–696 nm). Scale bars indicate 50 μm.
