## Supplementary tables for "Microbial mat compositions and localization patterns explain the virulence of black band disease in corals"

**Suppl. table. S1 Number of OTUs and the relative abundance in representative bacterial families.**

| | Family | Number of OTUs | Average of total relative abundance (%) | Ranging among the samples (%) |
|---|---|---|---|---|
| *Cyanobacteria* | *Desertifilaceae* | 7 | 42.57 | 1.39 – 77.68 |
| | *Oscillatoriaceae* | 3 | 1.10 | 0 – 9.75 |
| *Alfaproteobacteria* | *Rhodobacteraceae* | 44 | 5.64 | 0.04 – 26.03 |
| *Gammaproteobacteria* | *Vibrionaceae* | 12 | 4.01 | 0.15 – 18.79 |
| | *Alteromonadaceae* | 17 | 5.24 | 0.30 – 24.41 |
| | *Alteromonadaceae*_unclassified | 3 | 3.97 | 0.12 – 15.56 |
| | *Colwelliaceae* | 8 | 2.79 | 0.03 – 7.05 |
| | *Nitrincolaceae* | 7 | 2.23 | 0 – 13.19 |
| | *Pseudoalteromonadaceae* | 5 | 2.18 | 0.01 – 21.43 |
| | *Saccharospirillaceae* | 4 | 1.42 | 0.06 – 6.91 |
| *Deltaproteobacteria* | *Desulfovibrionaceae* | 8 | 3.14 | 0 – 13.41 |
| | *Desulfobacteraceae* | 7 | 2.52 | 0 – 8.72 |
| | *Desulfococcaceae* | 1 | 0.53 | 0 – 3.78 |
| *Campilobacterota* | *Arcobacteraceae* | 14 | 5.55 | 0.02 – 17.63 |
| | Rs-M59_termite_group | 1 | 0.51 | 0 – 6.11 |
| *Firmicutes* | *Caminicellaceae* | 1 | 1.20 | 0 – 3.99 |
| | *Lachnospiraceae* | 4 | 1.17 | 0 – 3.15 |
| | *Lachnospirales*_unclassified | 4 | 1.11 | 0 – 2.86 |
| | *Clostridiaceae* | 3 | 0.68 | 0 – 3.05 |
| *Bacteroidota* | *Flavobacteriaceae* | 67 | 1.06 | 0.05 – 5.79 |
| | *Saprospiraceae* | 76 | 0.83 | 0 – 4.00 |
| | *Bacteroidia*_unclassified | 49 | 0.53 | 0.02 – 1.72 |
| *Deferribacterota* | *Deferribacteraceae* | 1 | 0.61 | 0 – 3.78 |
| *Verrucomicrobiota* | P.palmC41_fa | 2 | 0.55 | 0 – 3.63 |
| Others (< 0.5%) | – | 1339 | 7.21 | 1.23 – 40.05 |

**Suppl. Table S2 Summary of representative OTUs (showing the total relative abundance with > 1%) in BBD with various linear-migration rates (n=12).**

| Family | OTU | The relative abundance ranged in proportion of samples (%) *[1] | Taxonomy *[2] | Closest sequence(s) in phylogenetic position *[3] | | | |
|---|---|---|---|---|---|---|---|
| | | | | Similarity (%) | GeneBank Acc. No. | Source | Reference |
| *Desertifilaceae* | OTU 1 | 1.38 – 77.67 (n=12) | *Roseofilum* AO1-A | 100 | NR_116573 | BBD in the Red Sea | (Rasoulouniriana et al. 2009) |
| | | | | 100 | KU579375.1 | BBD in Australia | (Buerger et al. 2016) |
| | | | | 100 | MH341659.1 | BBD in the Red Sea | (Hadaidi et al. 2018) |
| | | | | 100 | LC368145.1 | BBD in Japan | (Hutabarat et al. 2018) |
| *Oscillatoriaceae* | OTU 23 | 0.03 – 9.71 (n=3) | Uncultured bacteria | 98.01 | MT321585.1 | Boat launch in USA | (Berthold et al. 2021) |
| | | | | 96.02 | DQ446127.2 | BBD in Bahamas | (Sekar et al. 2008) |
| *Desulfovibrionaceae* | OTU 12 | 0.01 – 4.10 (n=10) | *Desulfovibrio* | 100 | AY497300.1 | BBD in Netherlands Antilles | (Frias-Lopez et al. 2004) |
| *Desulfobacteraceae* | OTU 5 | 0.02 – 7.98 (n=10) | *Desulfocella* | 100 | GU319466.1 | Coral treated in high pH from the Red Sea | (Meron et al. 2011) |
| | | | | 99.6 | KC527305.1 | Coral white plague disease in Thailand | (Roder et al. 2014) |
| *Rhodobacteraceae* | OTU 6 | 0.01 – 13.47 (n=11) | *Ruegeria* | 100 | MW589660.1 | Sea cucumber from Mexico | Quintanilla-Mena et al 2021, Unpublished |
| | | | | 100 | AF473938.1 | BBD in US Virgin Islands | (Cooney et al. 2002) |
| | OTU 24 | 0.01 – 6.96 (n=12) | *Thalassobius* | 100 | GU472129.1 | BBD in the Red Sea | Arotsker et al. 2010, Unpublished |
| | | | | 100 | MH341656.1 | BBD in the Red Sea | (Hadaidi et al. 2018) |

| | | | | | | | |
|---|---|---|---|---|---|---|---|
| *Arcobacteraceae* | OTU 8 | 0.54 – 10.84 (n=6) | Uncultured bacteria | 100 | MH341640.1 | BBD in the Red Sea | (Hadaidi et al. 2018) |
| | | | | 100 | EF123613.1 | BBD in US Virgin Islands | (Sekar et al. 2008) |
| | OTU 9 | 0.03 – 10.81 (n=9) | *Arcobacteraceae*_unclassified | 100 | MH341647.1 | BBD in the Red Sea | (Hadaidi et al. 2018) |
| | | | | 100 | HM768558.1 | BBD in Netherlands Antilles | (Klaus et al. 2011) |
| | OTU 11 | 0.01 – 5.46 (n=12) | *Arcobacteraceae*_unclassified | 98.88 | MH341652.1 | BBD in the Red Sea | (Hadaidi et al. 2018) |
| | | | | 98.88 | LT904749.1 | – | Latif-Eugenin et al. 2017, Unpublished |
| *Alteromonadaceae* | OTU 3 | 0.26 – 20.28 (n=12) | *Alteromonas* | 100 | LR812090.1 | – | Duchaud 2020, Unpublished |
| *Vibrionaceae* | OTU 7 | 0.01 – 18.74 (n=12) | *Vibrionaceae*_unclassified | 99.60 | CP045350.1 | Tank containing coral in Germany | Rockert et al 2019, Unpublished |
| | | | | 99.60 | NW828438.1 | Coral in Australia | Kuek et al. 2021, Unpublished |
| | | | | 99.60 | NW872696.1 | Coral | Loughran et al 2021, Unpublished |
| *Alteromonadales*_unclassified | OTU 2 | 0.12 – 15.56 (n=12) | *Alteromonadales*_unclassified | 99.60 | FJ202858.1 | Coral white plague in Puerto Rico | (Sunagawa et al. 2009) |
| *Colwelliaceae* | OTU 4 | 0.02 – 6.90 (n=12) | *Thalassotalea* | 100 | HM768682.1 | BBD in Netherlands Antilles | (Klaus et al. 2011) |
| | | | | 100 | KC527285.1 | Coral white plague disease in Thailand | (Roder et al. 2014) |
| *Nitrincolaceae* | OTU 14 | 1.13 – 13.19 (n=2) | *Marinobacterium* | 100 | MH341660.1 | BBD in the Red Sea | (Hadaidi et al. 2018) |
| | | | | 100 | FJ202983.1 | Coral white plague in Puerto Rico | (Sunagawa et al. 2009) |

| | | | | | | | |
|---|---|---|---|---|---|---|---|
| *Pseudoalteromonadaceae* | OTU 10 | 0.01 – 15.86 (n=11) | *Algicola* | 100 | FJ202088.1 | Coral white plague in Puerto Rico | (Sunagawa et al. 2009) |
| *Saccharospirillaceae* | OTU 19 | 0.01 – 5.14 (n=11) | *Thalassolituus* | 99.60 | HQ317342.1 | Biofilm of coral reef in Indonesia | Catalano er al 2010, Unpublished |
| | | | | 99.60 | MF039943.1 | Sponge | Keren et al 2010, Unpublished |
| *Caminicellaceae* | OTU 13 | 0.08 – 3.99 (n=9) | *Paramaledivibacter* | 100 | AY148309.1 | BBD in Australia | Cooney and Bythell 2002, Unpublished |
| | | | | 100 | AY348731.1 | Healthy tissue in coral disease from Australia | (Jones et al. 2004) |
| *Lachnospiraceae* | OTU 15 | 0.26 – 3.15 (n=9) | *Lachnospiraceae*_unclassified | 100 | MH341658.1 | BBD in Red Sea | (Hadaidi et al. 2018) |
| | | | | 100 | HM768582.1 | BBD in Netherlands Antilles | (Klaus et al. 2011) |
| *Lachnospirales*_unclassified | OTU 16 | 0.47 – 2.78 (n=9) | Lachnospirales_unclassified | 100 | GU471984.1 | BBD in Red Sea | Arotsker et al, 2020, Unpublished |
| | | | | 100 | GQ455295.1 | BBD in Red Sea | (Ben-Dov et al. 2011) |

*1 The proportions represent only the relative abundance of > 0.01% found in each sample.

*2 The Taxonomy defined base on Silva SSU ref v138.

*3 The closest sequences were chosen by high similarity and linage position in blastn (nucleotide collection nr/nt) and the function “Blast Tree View (Tree method: Fast Minimum Evolution and Max Seq Differences: 0.75)” in NCBI. When results hit multiple sequences, we selected those from sources related to BBD, coral disease, and healthy corals.

**Suppl. table S3 Partial correlation between representative bacterial families and liner-migration rates calculated by clr transformation matrixes.**

| | | *Spearman's rank correlation* | | |
|---|---|---|---|---|
| | Representative family | s | Correlation coefficient (rho) | *p* value*[1] |
| *Cyanobacteria* | *Desertifilaceae* | 130 | 0.5454545 | 0.07068 |
| | *Oscillatoriaceae* | 338 | –0.1818182 | 0.573 |
| *Alfaproteobacteria* | *Rhodobacteraceae* | 456 | –0.5944056 | **0.04575*** |
| *Gammaproteobacteria* | *Vibrionaceae* | 224 | 0.2167832 | 0.4991 |
| | *Alteromonadaceae* | 312 | –0.09090909 | 0.7832 |
| | *Alteromonadaceae*_unclassified | 212 | 0.2587413 | 0.4169 |
| | *Colwelliaceae* | 334 | –0.1678322 | 0.6037 |
| | *Nitrincolaceae* | 372 | –0.3006993 | 0.3425 |
| | *Pseudoalteromonadaceae* | 228 | 0.2027972 | 0.5281 |
| | *Saccharospirillaceae* | 212 | 0.2587413 | 0.4169 |
| *Deltaproteobacteria* | *Desulfovibrionaceae* | 194 | 0.3216783 | 0.3083 |
| | *Desulfobacteraceae* | 138 | 0.5174825 | 0.08865 |
| | *Desulfococcaceae* | 148 | 0.4825175 | 0.1154 |
| *Campilobacterota* | *Arcobacteraceae* | 42 | 0.8531469 | **0.0007719**** |
| | Rs-M59_termite_group | 188 | 0.3426573 | 0.2762 |
| *Firmicutes* | *Caminicellaceae* | 170 | 0.4055944 | 0.1926 |
| | *Lachnospiraceae* | 154 | 0.4615385 | 0.1338 |
| | *Lachnospirales*_unclassified | 150 | 0.4755245 | 0.1213 |
| | *Clostridiaceae* | 172 | 0.3986014 | 0.201 |
| *Bacteroidota* | *Flavobacteriaceae* | 418 | –0.4615385 | 0.1338 |
| | *Saprospiraceae* | 286 | 0 | 1 |
| | *Bacteroidia*_unclassified | 430 | –0.5034965 | 0.09875 |
| *Deferribacterota* | *Deferribacteraceae* | 142 | 0.5034965 | 0.09875 |
| *Verrucomicrobiota* | P.palmC41_fa | 302 | –0.05594406 | 0.869 |

*1 Significant of the *p* value are marked in bold with * $p < 0.05$ and ** $p < 0.01$.

**Suppl. table S4 Primers for bacterial community analysis and probes for FISH used in this study.**

| Analysis | Name | Sequence (5 '–3 ') | Targeted organism | *E.coli* position | Ref. |
|---|---|---|---|---|---|
| Bacterial community analysis | 515F | GTGCCAGCMGCCGCGGTAA | Most bacteria | 515 | (Aprill et al. 2015; Walters et al. 2016) |
| | 806R | GGACTACHVGGGTWTCTAAT | Most bacteria | 806 | (Aprill et al. 2015; Walters et al. 2016) |
| FISH | EUB338mix | GCTGCCTCCCGTAGG AGT<br>GCAGCCACCCGTAGGTGT<br>GCTGCCACCCGTAGGTGT | Most bacteria | 338 | (Amann et al. 1990; Daims et al. 1999) |
| | Arc94 | TGCGCCACTTAGCTGACA | *Arcobacteraceae* | 94 | (Snaidr et al. 1997) |
| | Non338 | ACATCCTACGGGAGGC | Non-target | – | (Wallner et al. 1993) |

**Suppl. table S5 Raw reads of amplicon sequences in this study.**

| Sample ID *1 | Number of read |
|---|---|
| AF_01 | 175,334 |
| AF_02 | 128,384 |
| AF_03 | 151,293 |
| AF_04 | 126,044 |
| AF_09 | 113,541 |
| AF_10 | 134,315 |
| SF_02 | 145,487 |
| SF_04 | 133,751 |
| SF_05 | 133,937 |
| SF_06 | 139,962 |
| SF_07 | 130,622 |
| SF_08 | 143,521 |

*1 Sample ID indicate as 'AF' and 'SF' that collected from Aka Island and Sesoko Island, respectively.

**Refences for Suppl. tables**


Amann RI, Binder BJ, Olson RJ, Chisholm SW, Devereux R, Stahl D a. (1990) Combination of 16S rRNA-targeted oligonucleotide probes with flow cytometry for analyzing mixed microbial populations. Appl Environ Microbiol 56:1919–1925.

Apprill A, McNally S, Parsons R, Weber L (2015) Minor revision to V4 region SSU rRNA 806R gene primer greatly increases detection of SAR11 bacterioplankton. Aquat Microb Ecol 75:129–137. doi: 10.3354/ame01753

Ben-Dov E, Siboni N, Shapiro OH, Arotsker L, Kushmaro A (2011) Substitution by Inosine at the 3′ -Ultimate and Penultimate Positions of 16S rRNA Gene Universal Primers. Microb Ecol 61:1–6. doi: 10.1007/s00248-010-9718-2

Berthold DE, Lefler FW, Laughinghouse HD (2021) Untangling filamentous marine cyanobacterial diversity from the coast of South Florida with the description of Vermifilaceae fam. nov. and three new genera: Leptochromothrix gen. nov., Ophiophycus gen. nov., and Vermifilum gen. nov. Mol Phylogenet Evol 160:107010. doi: 10.1016/j.ympev.2020.107010

Buerger P, Alvarez-Roa C, Weynberg KD, Baekelandt S, van Oppen MJH (2016) Genetic, morphological and growth characterisation of a new *Roseofilum* strain (Oscillatoriales, Cyanobacteria) associated with coral black band disease. PeerJ 4:e2110. doi: 10.7717/peerj.2110

Cooney RP, Pantos O, Le Tissier MD a., Barer MR, O'Donnell AG, Bythell JC (2002) Characterization of the bacterial consortium associated with black band disease in coral using molecular microbiological techniques. Environ Microbiol 4:401–413.

Daims H, Brühl A, Amann R, Schleifer KH, Wagner M (1999) The domain-specific probe EUB338 is insufficient for the detection of all Bacteria: development and evaluation of a more comprehensive probe set. Syst Appl Microbiol 22:434–444. doi: 10.1016/S0723-2020(99)80053-8

Frias-Lopez J, Klaus JS, Bonheyo GT, Fouke BW (2004) Bacterial community associated with black band disease in corals. Appl Environ Microbiol 70:5955–5962. doi: 10.1128/AEM.70.10.5955

Hadaidi G, Ziegler M, Shore-Maggio A, Jensen T, Aeby G, Voolstra CR (2018) Ecological and molecular characterization of a coral black band disease outbreak in the Red Sea during a bleaching event. PeerJ. doi: 10.7717/peerj.5169

Hutabarat PUB, Nguyen XH, Suda S (2018) Black Band disease-related (BBD) cyanobacterium from Okinawan corals. J Appl Phycol 30:3197–3203. doi: 10.1007/s10811-018-1507-1

Jones RJ, Bowyer J, Hoegh-Guldberg O, Blackall LL (2004) Dynamics of a temperature-related coral disease outbreak. Mar Ecol Prog Ser 281:63–77.

Klaus JS, Janse I, Fouke BW (2011) Coral Black Band Disease Microbial Communities and Genotypic Variability of the Dominant Cyanobacteria (CD1C11). Bull Mar Sci 87:795–821. doi: 10.5343/bms.2010.1050

Meron D, Atias E, Iasur Kruh L, Elifantz H, Minz D, Fine M, Banin E (2011) The impact of reduced pH on the microbial community of the coral Acropora eurystoma. ISME J 5:51–60. doi: 10.1038/ismej.2010.102

Rasoulouniriana D, Siboni N, Ben-Dov E, Kramarsky-Winter E, Loya Y, Kushmaro A (2009) Pseudoscillatoria coralii gen. nov., sp. nov., a cyanobacterium associated with coral black band disease (BBD). Dis Aquat Organ 87:91–96. doi: 10.3354/dao02089

Roder C, Arif C, Bayer T, Aranda M, Daniels C, Shibl A, Chavanich S, Voolstra CR (2014) Bacterial profiling of White Plague Disease in a comparative coral species framework. ISME J 8:31–39. doi: 10.1038/ismej.2013.127

Sekar R, Kaczmarsky L, Richardson L (2008) Microbial community composition of black band disease on the coral host Siderastrea siderea from three regions of the wider Caribbean. Mar Ecol Prog Ser 362:85–98. doi: 10.3354/meps07496

Snaidr J, Amann R, Huber I, Ludwig W, Schleifer KH (1997) Phylogenetic analysis and in situ identification of bacteria in activated sludge. Appl Environ Microbiol 63:2884–2896.

Sunagawa S, DeSantis TZ, Piceno YM, Brodie EL, DeSalvo MK, Voolstra CR, Weil E, Andersen GL, Medina M (2009) Bacterial diversity and White Plague Disease-associated community changes in the Caribbean coral Montastraea faveolata. ISME J 3:512–521. doi: 10.1038/ismej.2008.131

Wallner G, Amann R, Beisker W (1993) Optimizing fluorescent in situ hybridization with rRNA-targeted oligonucleotide probes for flow cytometric identification of microorganisms. Cytometry 14:136–143. doi: 10.1002/cyto.990140205

Walters W, Hyde ER, Berg-Lyons D, Ackermann G, Humphrey G, Parada A, Gilbert JA, Jansson JK, Caporaso JG, Fuhrman JA, Apprill A, Knight R (2016) Improved Bacterial 16S rRNA Gene (V4 and V4-5) and Fungal Internal Transcribed Spacer Marker Gene Primers for Microbial Community Surveys. mSystems 1:e00009-15. doi: 10.1128/mSystems.00009-15
