## Appendix for "Microbial mat compositions and localization patterns explain the virulence of black band disease in corals"

At family level, overall relative abundance (with top 24 families) of bacterial communities presented two families in *Cyanobacteria*: *Desertifilaceae*, *Oscillatoriaceae*, one family in *Alfaproteobacteria*: *Rhodobacteraceae*, seven families in *Gammaproteobacteria*: *Vibrionaceae*, *Alteromonadaceae*, *Alteromonadaceae*_unclassified, *Colwelliaceae*, *Nitrincolaceae*, *Pseudoalteromonadaceae*, *Saccharospirillaceae*, three families in *Deltaproteobacteria*: *Desulfovibrionaceae*, *Desulfobacteraceae*, *Desulfococcaceae*, two families in *Campilobacterota*: *Arcobacteraceae*, Rs-M59_termite_group, four families in *Firmicutes*: *Caminicellaceae*, *Lachnospiraceae*, *Lachnospirales*_unclassified, *Clostridiaceae*, three families in *Bacteroidota*: *Flavobacteriaceae*, *Saprospiraceae*, *Bacteroidia*_unclassified, one family in *Deferribacterota*: *Deferribacteraceae*, and one family in *Verrucomicrobiota*: P.palmC41_fa (**Fig. 3b** and **Suppl. table S1**).

To confirm the presence of cyanobacteria, SRB, SOB (except for the *Rhodobacteraceae* and *Arcobacteraceae*), and other heterotrophic bacteria that indicate BBD-microbiome reported in previous studies, we focused on the representative OTUs (operational taxonomic unit[s]) that considering compromised >1% of total relative abundance (**Suppl. table S2**). The most dominant bacterial family was the cyanobacterial family *Desertifilaceae,* which was currently proposed as new family in order *Oscillatoriales*[1], accounting for 42.57% of total relative abundance (**Suppl. table S1**). In the *Desertifilaceae*, OTU 1 contributed predominantly or rarely to the all microbial communities (in the ranging from 1.38 to 77.67%) and affiliated with *Roseofilum* AO1-A which was isolated from BBD in Australia[2] and detected across the Indo-Pacific (**Suppl. table S2**). As other dominant cyanobacteria affiliated with the family *Oscillatoriaceae,* OTU 23 was found in four samples from the both locations (in the ranging from 0.03 to 9.71%) with the linear-migration rates from 2.34 to 4.19 mm/day, and closely related with 98.01% similarity to *Vermifilum ionodolium* (accession No. MT321585.1) that retrieved from coastal limestone in USA. Interestingly, the second closest sequence (DQ446127.2) of OTU 23 according to 96.02% similarity were obtained from BBD in Bahamas (**Suppl. table S2**).

SRB belonging to the *Desulfovibrionaceae* and *Desulfobacteraceae*, respectively represented by OTU 12 and OTU 5, showed ranged in proportions between 0.01 and 4.10% and between 0.02 and 7.98%. The OTU 12 affiliated to the *Desulfovibrionaceae* had a 100% similarity with an uncultured delta proteobacterium clone CD22D5 (AY497300.1) that was detected in BBD mat in the Caribbean Sea (**Suppl. table S2**). The OTU 5 in *Desulfobacteraceae* was also closely related to uncultured bacterium Thai19_G02 (99.6% similarity), obtained from the coral disease white plague in Thailand, as second closest lineage (**Suppl. table S2**).

Besides the *Rhodobacteraceae* and *Arcobacteraceae,* we found only few reads of a common SOB member *Beggiatoa* spp. in our samples. In the family *Beggiatoaceae*, three OTUs (OTU 65,

OTU 140 and OTU 815) accounted for the relative abundance ranging from 0.0018 to 0.98% in seven samples with the linear-migration rates from 2.13 to 4.85 mm/day (data not shown).

In other heterotrophic bacteria, the represented OTUs in most of families such as *Vibrionaceae, Alteromonadales*_unclassified, *Colwelliaceae, Nitrincolaceae, Pseudoalteromonadaceae, Caminicellaceae, Lachnospiraceae,* and *Lachnospirales*_unclassified showing varied relative abundances among our samples were linked with BBD and other coral diseases (**Suppl. table S2**). Notably, OTU 7 in *Vibrionaceae* showed high relative abundance with 18.74% in samples with the fastest migration rates (**Fig. 3** and **Suppl. table S2**). Although the amplicon sequences could only provide good resolution at the genus level[3], the best hit list of the sequences indeed included several *Vibrio* type strains in the blast search.
